# Brain-heart coupling shapes large scale brain dynamics

**DOI:** 10.1101/2025.07.14.664815

**Authors:** Saurabh Sonkusare, Kartik Iyer, Luke Hearne, Richa Phogat, Johan van der Meer, Sasha Dionisio, James M. Shine, Michael Breakspear

## Abstract

Functional brain networks reconfigure to support adaptive behaviour, balancing integration and segregation. These dynamics are influenced by neuromodulatory arousal, reflected in heart rate (HR) fluctuations. We leverage intracranial EEG (iEEG), functional MRI (fMRI), and HR recordings during emotional movie viewing to probe neurocardiac contributions to large-scale network dynamics. Network dynamics using high-frequency iEEG activity (60– 140 Hz), an established marker of neuronal firing, reveal that greater network integration tracks elevated HR, while segregation aligns with lower HR. However, we observe an inverted relationship between HR and network dynamics in a separate fMRI dataset utilising the same movie, and replicate this in another independent fMRI dataset. Biophysical modelling of BOLD from iEEG reproduces the iEEG-HR findings, suggesting neurovascular transformation may obscure neural-cardiac links in fMRI. These results link large-scale network dynamics to physiological arousal through HR driven adrenergic-cholinergic neuromodulation and highlight the complex interplay between neural activity, autonomic regulation, and haemodynamics.

## Introduction

The brain’s ability to adapt to changing internal bodily states and the external environmental milieu depends on the dynamic reconfiguration of its functional networks, switching between localized segregated processing and global information integration^1–3^. While integration facilitates communication between distinct systems, segregation preserves the autonomy of specialized subnetworks for parallel processing and functional specialization^4,5^. The dynamic trade-off between functional segregation and integration underlies adaptive behaviour and is increasingly viewed as a core organizing principle of large-scale neural dynamics across species and behavioural states^6^.

The shifts in the balance between integration and segregation are not static properties of the brain but vary across time and behavioural context, closely tracking changes in arousal and physiological state^7,8^. Arousal, a global physiological state governing vigilance, attention, and emotional responsiveness, plays a critical role in modulating large-scale network architecture ^9^. Arousal is predominantly mediated by brainstem-originating neuromodulatory systems, including noradrenergic and cholinergic pathways, which exert widespread influence over cortical excitability and synaptic gain, dynamically adjusting network topology in response to internal and external demands^10,11^. Noradrenaline enhances neuronal excitability and is believed to facilitate large-scale network integration, promoting adaptive responses to salient or unexpected stimuli^12^. In contrast, acetylcholine sharpens sensory precision and aids in functional segregation, supporting fine-grained, context-specific processing^13^. These neuromodulatory influences govern the balance between integration and segregation, linking arousal to large-scale neural reconfiguration.

Despite their central role in regulating network dynamics, direct in vivo measurement of neuromodulatory activity remains technically challenging due to the diffuse and transient nature of neurotransmitter signalling^14^. Consequently, indirect physiological markers provide a valuable alternative for assessing neuromodulatory influences on brain function. Heart rate (HR), controlled by the autonomic nervous system, reflects the interplay between adrenergic (sympathetic) and cholinergic (parasympathetic) activity and serves as a reliable index of central arousal-regulating mechanisms^15,16^. Elevated HR is associated with sympathetic dominance, reflecting heightened adrenergic drive that enhances alertness and cognitive resource allocation^17^. Conversely, HR deceleration is linked to parasympathetic (vagal) activity, indicative of cholinergic modulation that promotes localized processing and attentional stability^17^. These autonomic HR fluctuations provide an accessible physiological readout of neuromodulatory balance and its influence on large-scale neural dynamics and cognition^18^. However, direct empirical evidence linking neurophysiological signals, network topology, and autonomic state remains sparse.

A key hypothesis underlying these HR-driven neuromodulatory effects centres on the complementary roles of two brainstem nuclei: the noradrenergic locus coeruleus (LC) and the cholinergic nucleus basalis of Meynert (NBM)^19,20^ *(Fig. 1A).* The LC, through its widespread projections, is thought to facilitate global integration by enhancing excitability and coordinating adaptive responses to salient or unexpected stimuli, particularly under heightened arousal^21^. In contrast, the NBM is implicated in promoting functional segregation by sharpening sensory precision and modulating local processing, with acetylcholine increasing signal fidelity in cortical circuits^13,22^. These opposing neuromodulatory influences dynamically shape network topology, linking arousal-mediated shifts in brain state to large-scale neural reorganization^23^.

**Figure 1.**
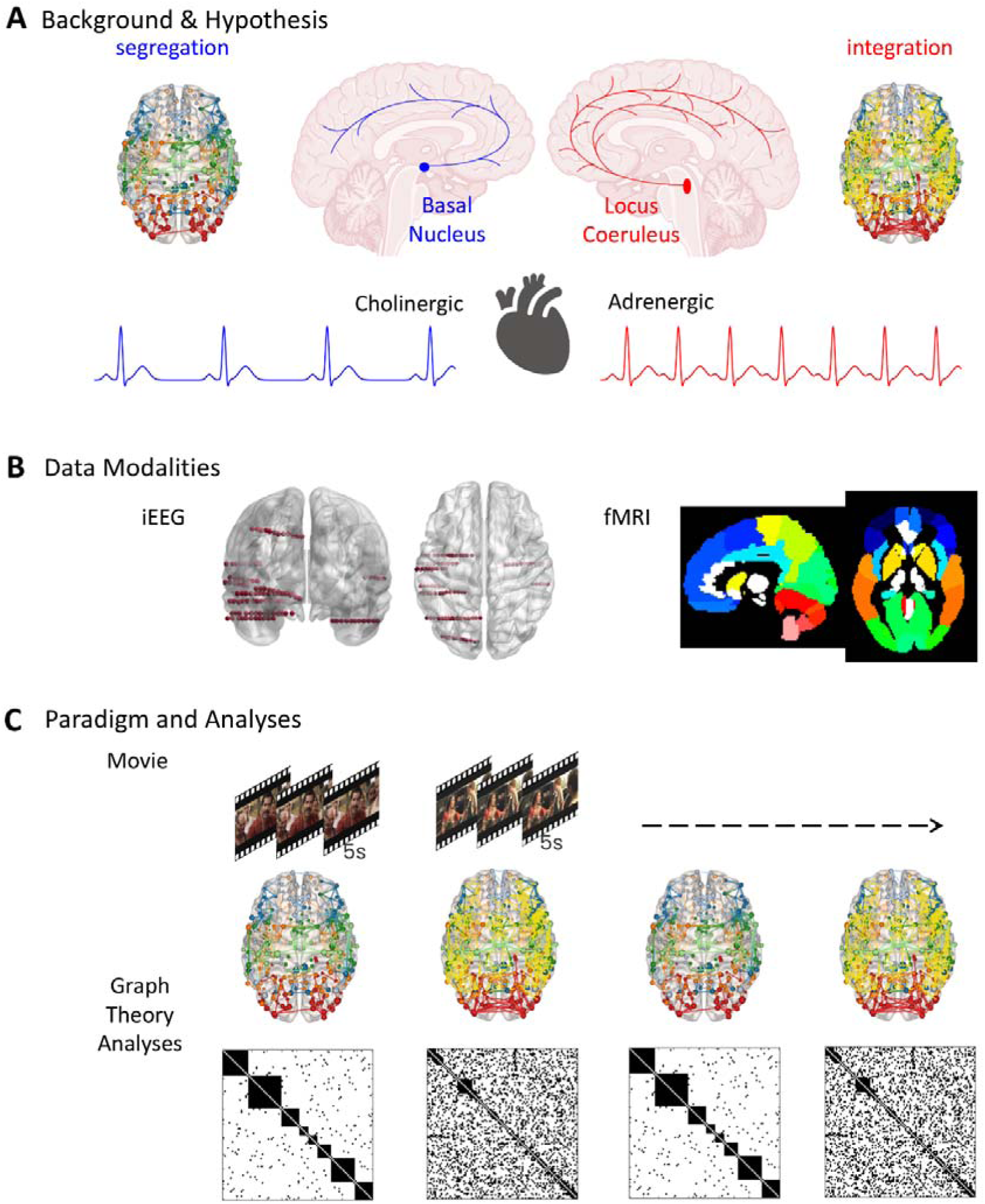
Hypothesis and schema for analytical approach. **(A)** The Locus Coeruleus, a noradrenergic brainstem nucleus, has diffuse projections throughout the cortex and subcortex, facilitating the integration of brain networks. In contrast, the Basal Nucleus (of Meynert), a cholinergic brainstem nucleus, exhibits sparse projections to localized networks, promoting segregation in brain networks. While in-situ neurotransmitter dynamics are challenging to measure directly, heart rate (HR)—controlled by the adrenergic-cholinergic balance—serves as a proxy for assessing this balance. **(B)** To investigate our hypothesis, we utilized intracranial EEG (iEEG) data, which offers millisecond temporal resolution at a meso-network spatial level, and functional MRI, which provides macroscale global brain coverage. **(C)** A 20-minute emotional movie was used to probe dynamic graph properties, with integration and segregation metrics extracted from windowed data (sliding window analyses for fMRI data) to generate their respective time series. Mean HR was computed for each corresponding time window to establish a link with the integration and segregation time series.

To investigate how arousal-related physiological signals influence brain network dynamics, we leveraged multimodal datasets of intracranial EEG (iEEG) and functional MRI (fMRI), with simultaneously acquired HR, utilising time-resolved graph theoretical analyses *(Fi.g 1 B,C).* Specifically, we used iEEG to investigate the link between functional network reconfiguration and HR fluctuations during naturalistic movie viewing—a paradigm that evokes dynamic shifts in arousal and affective engagement. iEEG provides direct access to mesoscale neural activity with millisecond precision, allowing for the examination of fast network transitions. We focus on high-frequency broadband (HFB) activity (60–140 Hz), a well-established marker of local neuronal firing rates^24,25^ to derive time-resolved measures of network topology. By correlating fluctuations in integration and segregation with continuous HR recordings, we test whether functional network reconfiguration tracks the dynamic balance between sympathetic (adrenergic) and parasympathetic (cholinergic) influences.

Recent evidence suggests that the relationship between neural activity, hemodynamic signals, and physiological state is non-linear and context-dependent, raising the possibility that BOLD-based measures of integration and segregation may not fully recapitulate patterns derived from direct neural recordings^26,27^. Consequently, we further examine whether similar brain-heart coupling patterns emerge in functional MRI (fMRI) data from a separate group of healthy participants viewing the same movie. fMRI, while lacking the temporal precision of iEEG, offers the opportunity to assess large-scale functional dynamics across the whole brain, allowing for cross-modal validation of the observed brain-heart coupling patterns. To validate our results from iEEG and fMRI, we employ a Balloon-Windkessel hemodynamic model to simulate BOLD signals from the iEEG data, allowing us to directly test whether putative discrepancies between modalities reflect true differences in underlying neural processes, or instead arise from differences in the neural-hemodynamic transformation itself.

## Results

### iEEG and fMRI datasets during naturalistic movie viewing

We analyzed iEEG and a separate fMRI dataset in which participants had viewed an unedited 20-minute movie previously shown to elicit robust intersubject correlations (ISC) and affective engagement^28,29^. We first localised the iEEG electrode contacts in gray matter (Fig 2 A). Local field potentials from iEEG carry complex information in various frequencies; however, the high-frequency broadband (HFB) activity has been demonstrated to be a robust marker of neuronal firing rates ^24^ and is a direct correlate of the BOLD signals^30^. We thus focussed on the HFB activity (60–140 Hz) (see methods). We computed HFB activity for the whole movie data and segmented it into 5-second non-overlapping windows, ensuring sufficient heartbeats per segment for robust mean heart rate (HR) estimation. Connectivity matrices were derived for each segment using Pearson’s correlation between HFB activity, thresholded at 0.2 to eliminate spurious connections. We chose widely used graph measures of segregation (modularity) and integration (global efficiency), which were calculated from these connectivity matrices, generating time series of integration and segregation for each patient. Simultaneous ECG recordings provided mean HR for each 5-second segment. Patient-specific Pearson correlations between integration-HR and segregation-HR dynamics were averaged to obtain group-level correlations. Statistical significance was assessed using permutation testing (5000 iterations) to generate null distributions.

**Figure 2.**
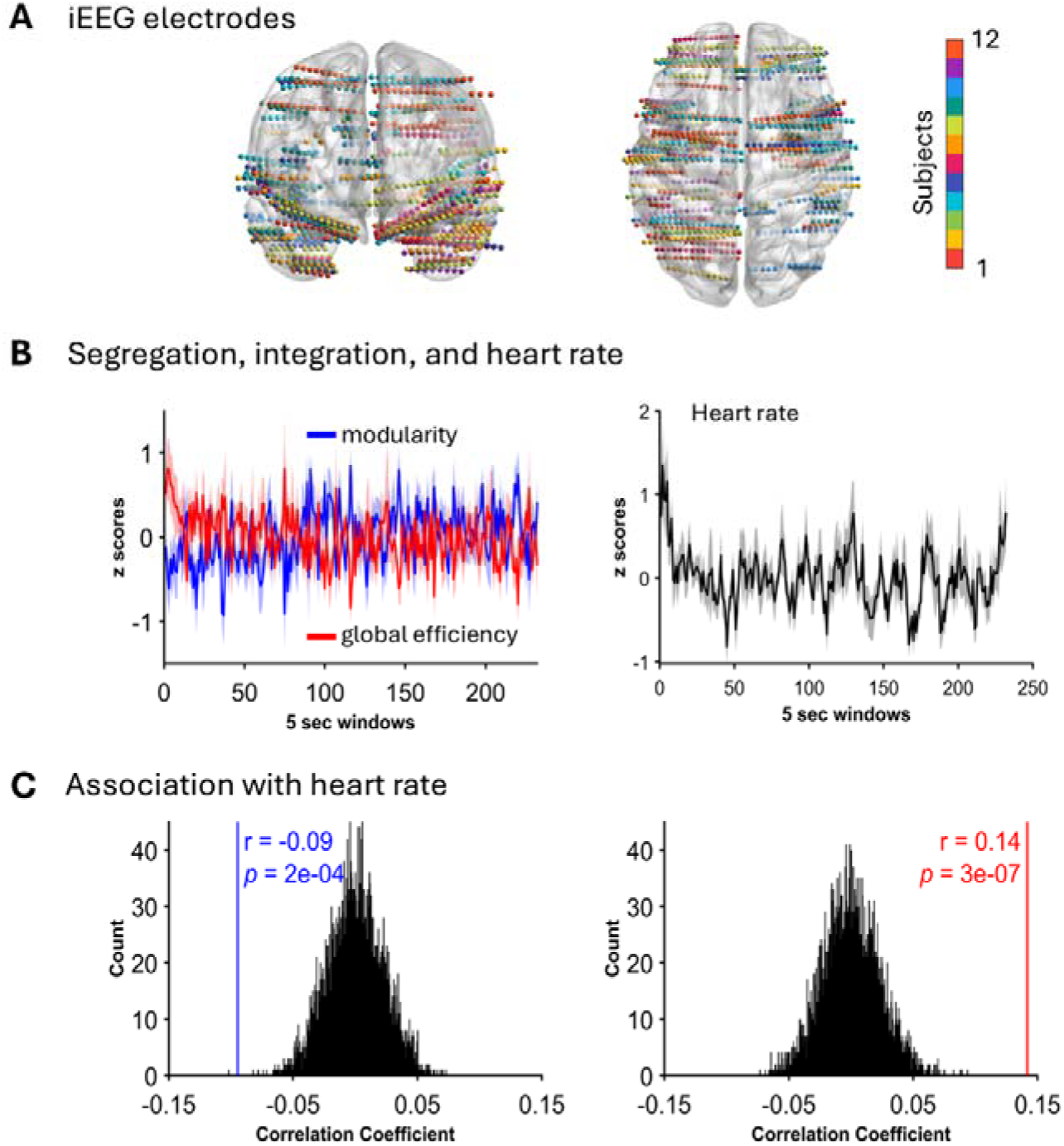
iEEG analysis and results. **(A)** The iEEG electrodes from all the patients were transformed into MNI space. All the electrode and their contacts are shown. The analyses were undertaken for the contacts in the gray matter. **(B)** The mean time series for graph metrics of integration and segregation are shown (left) along with the HR (right). Shading indicate SEM across participants. **(C)** Null distribution obtained with 5000 permutations showing statistical significance of negative correlation between segregation and HR whilst a positive correlation was found between integration and HR.

The fMRI dataset included 17 healthy participants watching the same movie. fMRI analyses followed the same approach as for iEEG, except sliding window analyses was employed for dynamic analyses of fMRI signals (window ∼25 s, sliding step 6.6 s, replicated at 44s) (see methods)^8,31^.

### HR links to iEEG integration and segregation dynamics

Group-averaged integration and segregation time series along with heart rate time series are shown in (*Fig. 2B*). The statistical significance of the link between integration-segregation dynamics and HR was tested with permutation testing (see Methods). This revealed a significant negative correlation between segregation and HR (r = −0.09, p < 0.001) and a significant positive correlation between integration and HR (r = 0.14, p < 0.001) (*Fig. 2C*).

### Association with movie features and iEEG dynamics

We conducted Pearson’s correlation analyses to examine the relationships between the grouped mean time series of integration and segregation, HR, and movie features, including portrayed facial emotions, visual saliency, audio, and luminance (*Fig. 3*). After correcting for multiple comparisons using the false discovery rate (FDR), we observed several significant associations. As expected, integration dynamics showed a positive correlation with HR (r = 0.16, p_FDR_ < 0.05). Only integration dynamics were strongly positively correlated with the luminance of the movie (r = 0.23, p_FDR_ < 0.05). Additionally, audio amplitude was positively correlated with HR (r = 0.20, p_FDR_ < 0.05) and with portrayed facial emotions (r = 0.25, p_FDR_ < 0.05). Conversely, luminance exhibited a negative correlation with portrayed emotions (r = −0.18, p_FDR_ < 0.05) and a positive correlation with visual saliency (r = 0.19, p_FDR_ < 0.05).

**Figure 3.**
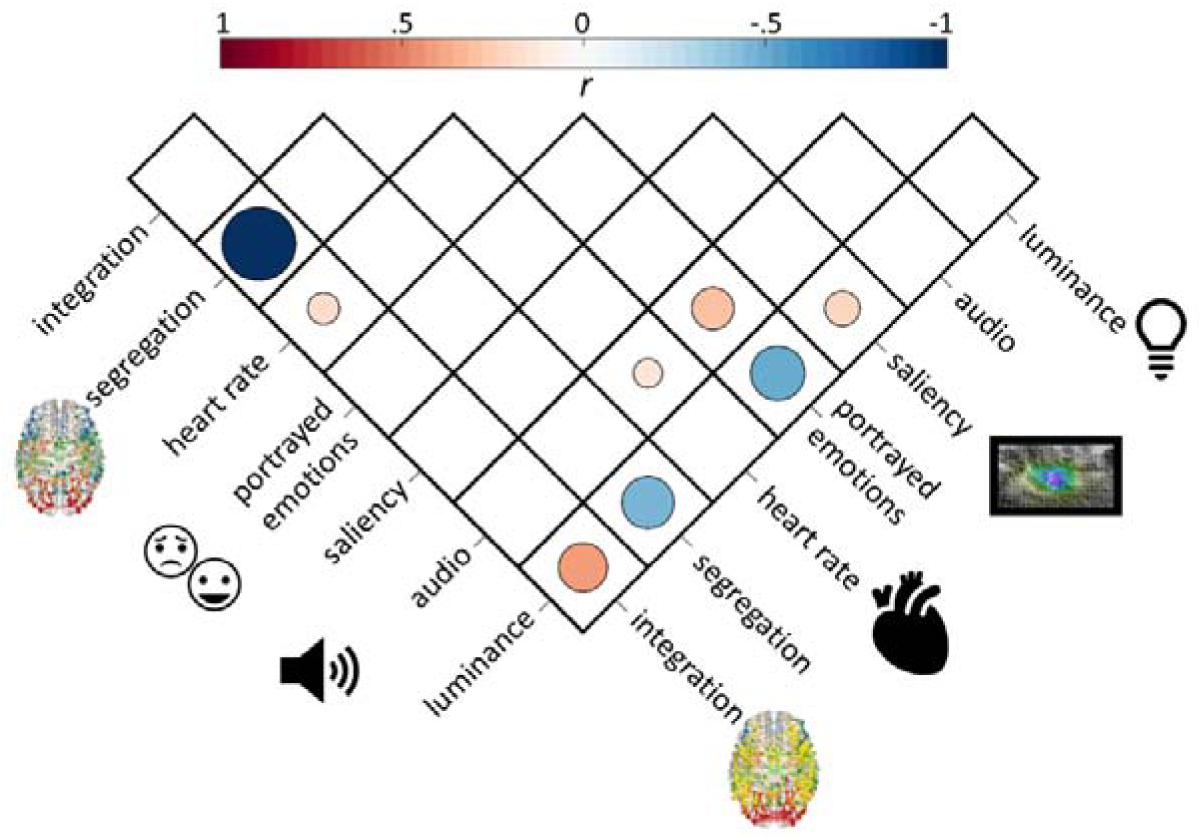
Association with movie features. Pearson’s correlation coefficient values (*r*) between integration, segregation, HR and movie features of portrayed emotions, salience (visual), audio; and luminance. Correlation coefficient values that were significant after multiple comparisons correction (p<.05) are shown.

### Opposing pattern of iEEG results with BOLD fMRI

We used an independent fMRI data, which used the same movie stimulus, to investigate whether iEEG graph properties and their link to HR extend to whole-brain BOLD dynamics. We first validated if the mean HR time series obtained from iEEG data showed similarity with that of healthy participants in the fMRI cohort. We found a strong positive correlation (r = .37,p = 2e-7) (*Fig. 4A*), indicating similar physiological arousal and engagement. Then, using the AAL atlas (a whole-brain atlas with cortical, subcortical, and cerebellar regions) ^32^, we extracted the region-wise BOLD signals. To capture time-resolved changes in integration and segregation, we employed a sliding-window approach. Given that heart rate fluctuations during movie viewing occur on shorter timescales than in resting-state paradigms, we prioritised a finer temporal resolution (window size = 11 TRs, 24.2 s; step = 3 TRs, 6.6s) to avoid oversmoothing transient effects. To ensure robustness, we replicated our findings using a conventional 44-second window ^8,33^, confirming that results were not driven by window-length selection (*see below*). In contrast to iEEG findings, fMRI analyses revealed the opposite pattern: HR was positively correlated with network segregation (r = 0.15, p < 0.001) and negatively correlated with integration (r = −0.17, p < 0.001) (*Fig. 4B*). We also replicated the results when all fMRI time series were shifted forward in time by 6.6s (3 TRs) to account for the approximate hemodynamic delay (*Supplementary Fig S1 A).* We conducted supplementary analyses by convolving HR time series with the empirically derived cardiac response function (CRF)^34^ (see Methods). Applying this convolution replicated our findings of a positive association of HR with segregation (r = 0.10, p =.003) and a negative association of HR with integration (r = −0.14, p = 0.00002).

**Figure 4.**
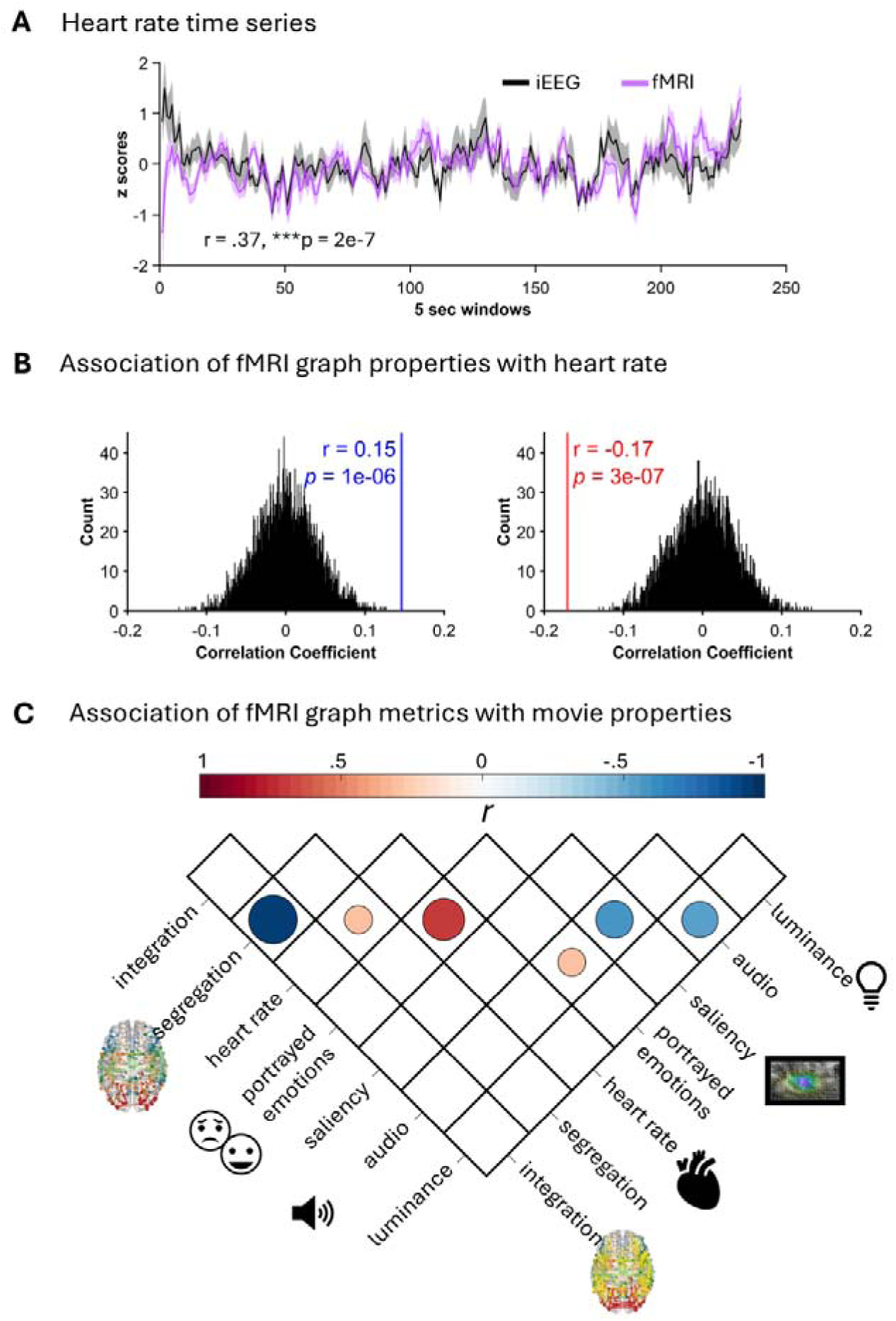
Consistent heart rate dynamics and fMRI results. **(A)** Heart rate time series from iEEG data and that from fMRI data (resampled to same 5-second segments) showed a significant positive correlation of the mean time series, indicating robust and similar physiological responses among patients and healthy participants for the same movie **(B)** Null distribution derived from 5,000 permutations, demonstrating the statistical significance of a positive correlation between segregation and HR and a negative correlation between integration and HR. **(C)** Pearson’s correlation coefficient values (*r*) between integration, segregation, HR and movie features of portrayed emotions, salience (visual), audio; and luminance. Correlation coefficient values that were significant after multiple comparisons correction (p<.05) are shown.

Furthermore, we replicate our findings when using longer time windows of 20 TRs (44 seconds) (*Supplementary Fig S1 B*). We also replicate our fMRI results when using a finer-scale brain atlas, specifically the Briannetome atlas, which has 274 parcels that include 210 cortical, 36 subcortical and 28 cerebellar regions ^35^ (*Supplementary Fig S2 A*). The results also hold true when using this atlas and employing a longer time window of 44 seconds (*Supplementary Fig S2 B*). In fMRI studies, while global efficiency is a widely used measure of network integration, the participation coefficient (PC) metric offers a complementary perspective by quantifying the extent of a node’s connectivity across different network modules ^3,36,37^. Hence, we also included PC to assess network topology in relation to heart rate (HR). Integration obtained via PC showed a significant negative correlation with HR (r = −0.10, p = 0.001; *Supplementary Fig. S3*), further supporting our findings of reduced integration with higher HR.

Finally, given that alpha activity (8-12 Hz) has been shown to be inversely correlated with BOLD signals^38,39^, we also undertook band-pass filtering of iEEG recordings in this alpha frequency range (8-12 Hz) and applied similar iEEG analyses as high-frequency activity. We found no significant association of alpha iEEG activity-derived integration segregation with HR (*Supplementary Fig S4)*.

### Association with movie features with fMRI dynamics

We conducted Pearson’s correlation analyses to examine the relationships between the grouped mean time series of integration and segregation derived from fMRI, HR, and movie features reported earlier (*Fig. 4C*). After correcting for multiple comparisons using the false discovery rate (FDR), we observed several significant associations. Segregation dynamics showed a strong negative correlation with integration (r = -.97, p_FDR_ < 0.05) and a positive correlation with HR (r = 0.32, p_FDR_ < 0.05). Portrayed emotions were strongly positively correlated with HR (r = 0.70, p_FDR_ < 0.05) and with audio (r = 0.31, p_FDR_ < 0.05). Audio exhibited a negative correlation with saliency (r = −0.58, p_FDR_ < 0.05) and with luminance (r = −0.54, p_FDR_ < 0.05).

### Replication of BOLD findings in an independent fMRI dataset

To assess the robustness of these effects, we analyzed an independent open-access fMRI dataset (N=30) with concurrent HR recordings acquired during movie viewing (see Methods). Keeping analytical window durations and sliding steps similar to our previous analyses, we used 26-second windows (20 TRs) with a 6.5 second sliding step (5TRs). These analyses confirmed the fMRI findings, demonstrating a positive correlation between segregation and HR (r = 0.11, p < 0.001) and a negative correlation between integration and HR (r = −0.16, p < 0.001) (*Fig. 5A*). These results demonstrated a reproducible association between autonomic activity and large-scale network organization in BOLD fMRI, distinct from the coupling observed in iEEG.

**Figure 5.**
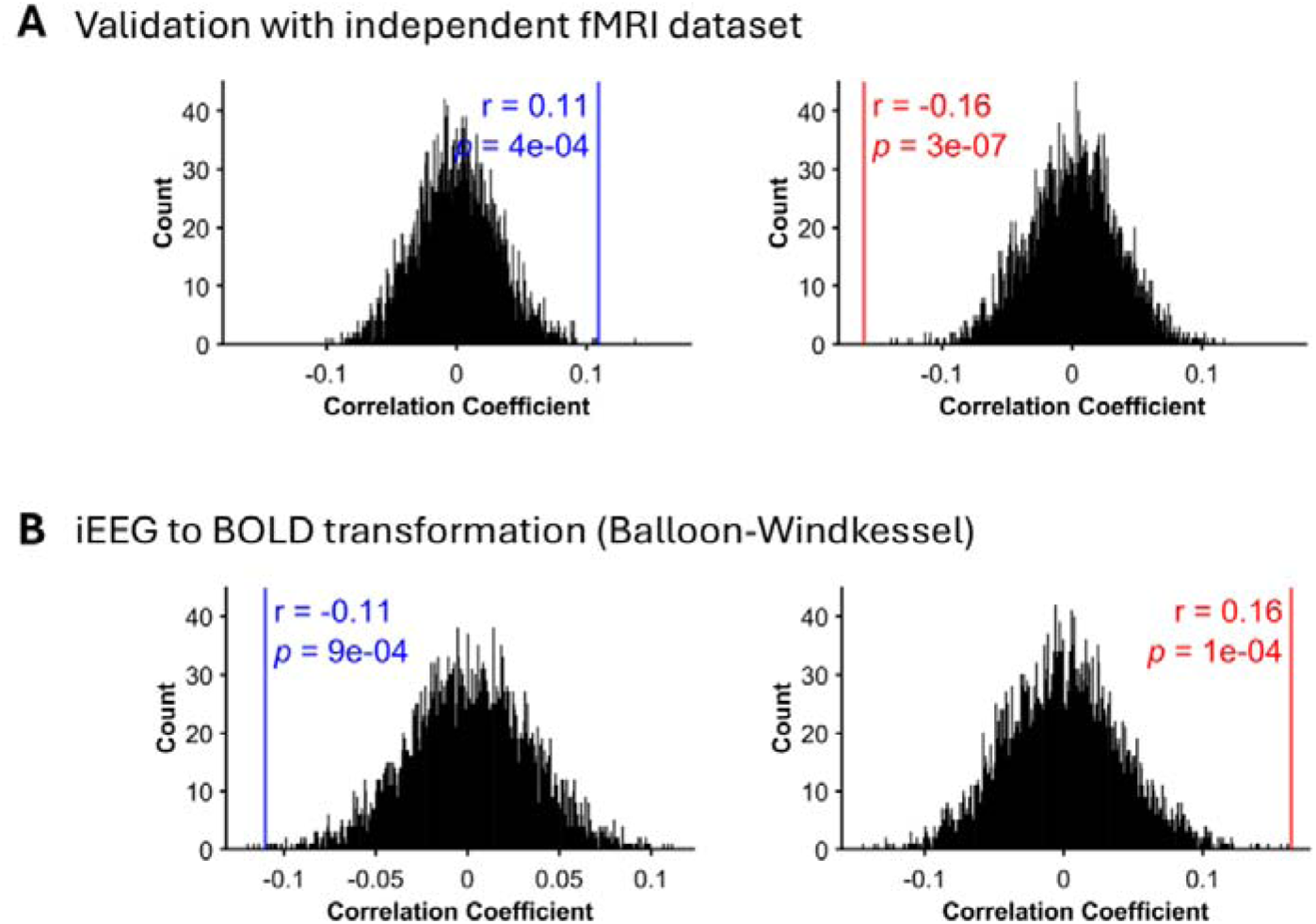
Replication with external fMRI dataset and modelling iEEG to BOLD transformation. **(A)** Replication of these findings in an independent fMRI dataset during movie viewing, confirming a positive correlation between segregation and HR and a negative correlation between integration and HR. **(B)** To bridge iEEG and fMRI findings, iEEG data were transformed into simulated BOLD signals using the Balloon-Windkessel model. The integration-segregation metrics computed from these signals were linked to HR, revealing results consistent with the original iEEG findings and not replicating the fMRI results.

### Reconciling iEEG–fMRI discrepancies via Balloon-Windkessel modelling

To resolve the divergence between iEEG and fMRI findings, we simulated BOLD responses from iEEG-derived neural activity using the Balloon-Windkessel model. We used identical window length and sliding steps as used in our first fMRI analyses i.e. window size of 11TRs and sliding steps of 5 TRs. The simulated BOLD signals did not replicate the empirical fMRI results but instead mirrored iEEG findings, showing a negative correlation between segregation and HR (r = −0.11, p < 0.001) and a positive correlation between integration and HR (r = 0.16, p < 0.001) (*Fig. 5B*).

## Discussion

Our findings demonstrate that large-scale brain network dynamics are tightly coupled to autonomic fluctuations, with heart rate reflecting underlying neuromodulatory balance. iEEG analyses revealed that increased network integration aligns with elevated heart rate, whereas greater segregation corresponds with lower heart rate. However, fMRI analyses from two independent datasets exhibited an opposing pattern. Transforming the iEEG signals into synthetic BOLD signals using the classic balloon model recapitulated iEEG-HR findings, highlighting complex, nonlinear interactions between neural activity, vascular dynamics, and autonomic regulation.

### Neurocardiac Coupling and Large-Scale Network Dynamics

Our iEEG findings demonstrate that fluctuations in autonomic state dynamically shape large-scale brain network organization. Periods of elevated HR were associated with increased network integration, consistent with the role of heightened arousal in facilitating global neural coordination and optimizing responsiveness to external stimuli. This pattern aligns with prior work showing that states of increased physiological activation promote widespread cortical communication, enabling rapid information transfer and adaptive cognitive flexibility^40^. In contrast, HR deceleration coincided with greater network segregation, suggesting that lower arousal states favour localized processing^41^. These findings provide direct electrophysiological evidence that shifts in physiological state are closely tied to transitions in functional brain topology, supporting the view that neural network dynamics are tightly coupled to autonomic regulation^15,18^. Recent work also suggests large-scale fMRI signal fluctuations are not random or artifactual but reflect coordinated neurophysiological events tied to the body’s autonomic arousal system^42^.

Our iEEG results also align with theoretical models implicating two key brainstem-originating neuromodulatory systems — the noradrenergic LC and the cholinergic NBM — posited to play complementary roles in governing network dynamics. Specifically, the LC, the brain’s primary source of adrenaline (noradrenaline), enhances cortical excitability and global integration, particularly during heightened arousal or novelty^43^. Rodent studies also link LC firing to sensory gain and behavioural arousal^19,44^. Some evidence of limited and low-frequency optogenetic stimulation of LC in rats did not uniformly modulate global brain activity^45^. In contrast, cholinergic projections from the NBM enhance functional segregation by sharpening sensory precision and promoting local processing during sustained attention^20^. The NBM modulates cortical desynchronization and local circuit dynamics, as shown by lesion and pharmacological studies in primates^46^. Computational modelling also suggests these network reconfigurations are mediated by ascending neuromodulation via their opposing influence on the sharpness of the gain function in local neural populations^11,23^.

### Divergent relationships across modalities: electrophysiology versus haemodynamics

A key finding of our study is the striking reversal of the HR–network relationship between iEEG and fMRI data. HFB activity directly reflects local neuronal activity^24^. Although the neural basis of BOLD signals is thought to derive largely from local HFB^30^, BOLD also integrates neural, vascular, and metabolic processes over longer timescales^47^. Notably, alpha oscillations (8-12 Hz) are inversely related to BOLD signals^38,39,48^. However, graph measures computed on alpha activity did not show significant associations with HR (*Supplementary Figure S3*). These divergent results highlight the profound influence of measurement modality on apparent relationships between network dynamics, arousal and neurovascular coupling.

Although fMRI pipelines sometimes include explicit correction for physiological “noise” signals such as heart rate and respiration^49^, our core findings persisted across preprocessing strategies, underscoring their robustness across analytical variations. Additionally, Balloon-Windkessel simulations failed to reproduce the empirical fMRI findings, instead replicating the iEEG-based findings despite the effective low-pass filtering effect of the HRF and the longer time windows employed to match the fMRI pipeline. This suggests that the opposing HR-network relationships cannot be explained by simple transformations of neural activity into BOLD signals. Instead, they likely reflect complex interactions between neuromodulatory tone, vascular reactivity, and systemic physiology^27^.

A possible physiological mechanism may explain the discordant results. Cholinergic-driven vasodilation via muscarinic receptors could enhance BOLD signals in segregated networks during low HR states. In contrast, adrenergic vasoconstriction (mediated by α-adrenergic receptors) may suppress BOLD signals in integrative networks during high HR states^50^. This opposing vascular regulation may explain why the HR-BOLD relationship inverts relative to iEEG’s direct neural measures. This mechanism aligns with evidence of strong negative correlations between HR and resting-state fMRI signals with peak effects delayed by 6–12 seconds^51^. Together, these findings reveal how autonomic-driven hemodynamic shifts can distort the coupling between neural activity and fMRI signals.

### Naturalistic paradigms for revealing neuromodulatory network dynamics

Unlike resting-state paradigms, which lack structured cognitive and affective engagement, movie viewing elicits dynamic fluctuations in arousal, cognitive processing and robust peripheral physiological changes^52–54^. However, despite using four movie features, we found that integration dynamics were only positively correlated with luminance of the movie frames. Higher luminance strengthens feedforward signals from the retina to V1 and higher-order regions and also promotes gamma-high gamma band oscillations, facilitating large-scale neural synchronization involved in global integration^55,56^. Converging evidence from mice studies supports this interpretation, showing that larger pupil diameters coincide with desynchronized, activated cortical states, whereas smaller pupils reflect more synchronized, low-arousal states^57^. Notably, integration dynamics were not significantly associated with visual saliency, implying saliency may represent a higher-order property^58^. The structured external drive of the movie likely minimized intersubject variability in network dynamics as well as heart rate, while resting-state fMRI, being an unconstrained state, induces vast intersubject variability^59^. Additionally, microsleeps prevalent during resting-state scans^59,60^ may further obscure large-scale network dynamics in relation to arousal, highlighting the necessity of task engagement when investigating neuromodulatory influences.

### Limitations and future directions

While our multimodal approach offers significant insights, several limitations must be acknowledged. First, the iEEG data were obtained from patients with focal epilepsy, which may restrict generalizability to healthy populations. Interictal spikes could also influence network properties by potentially exaggerating correlations. However, such spikes occur at different times across participants, mitigating time-locked effects. Methodological differences between modalities also warrant consideration. iEEG network properties were computed over shorter 5-second epochs to align with HR fluctuations, while fMRI analyses relied on sliding window connectivity estimates with relatively long windows (25-45 seconds) to ensure sufficient signal-to-noise ratios. These differences likely contribute to the observed divergence between modalities. Additionally, the nature of preprocessing steps for fMRI can introduce confounds and thus remains an important issue requiring deeper investigation. However, the replication of fMRI results in an independent dataset with different preprocessing pipelines suggests that these findings are not artifacts of specific processing steps but reflect robust phenomena. Our findings raise critical questions for the field regarding the relationship between BOLD signals, physiological measures, and brain-body coupling. Future work should further explore these methodological nuances to refine our understanding of neuromodulatory influences on brain network dynamics.

## Methods

### iEEG data and analyses

#### Participants

12 patients with medically intractable epilepsy, implanted with stereotactic electrodes for clinical evaluation at Mater Advanced Epilepsy Unit, Mater Hospital Brisbane, participated in this study. The choice of regions for electrode implantation was based on clinical criteria alone. Patient characteristics are provided in *Table 1*. Participants gave written informed consent to participate in the study and were free to withdraw from the study at any time. The study was approved by the Human Research Ethics Committees of the Mater Hospitals and the QIMR Berghofer Medical Research Institute and performed in agreement with the Declaration of Helsinki.

**Table 1.** Patient demographics.

| Id | Sex | Age (Y) | Handedness | Sz onset Age | Main diagnosed epileptogenic zone |
| --- | --- | --- | --- | --- | --- |
| P1 | M | 46 | R | 37 | Left mesial Orbitofrontal/gyrus rectus |
| P2 | M | 31 | R | 24 | Left temporal neocortical region and hippocampus |
| P3 | M | 40 | R | 28 | Focal left mesial temporal (hippocampus, amygdala, temporal pole) |
| P4 | F | 16 | R | 8 | Right amygdala/uncus |
| P5 | F | 22 | L | 15 | Amygdalo-perisylvian (Peri-rhinal/entorhinal cortex), Amygdala-hippocampal region |
| P6 | M | 56 | R | 17 | Right orbitofrontal cortex |
| P7 | F | 26 | R | 15 | Focal, right posterior middle and superior temporal gyrus; right supramarginal gyrus |
| P8 | M | 34 | R | 32 | Right supracalcarine, inferior cuneus |
| P9 | M | 41 | R |  | Left anterior to mid precuneus |
| P10 | M | 41 | R | 27 | Amygdala and entorhinal region |
| P11 | M | 21 | R | 6 | Focal, left parietal operculum |
| P12 | M | 38 | R | 12 | right planum polare; right temporal lobectomy had been performed prior |

#### Intracranial EEG (iEEG) recordings

Implantation of intracerebral electrodes with multiple contacts (DIXI, 10-15 contacts, electrode diameter: 1 mm, intercontact spacing 1.5 mm) was performed with a stereotactic procedure and planned individually based on the likely seizure onset zone inferred by the epilepsy clinician (author SD) from non-invasive pre-operative studies. sEEG signals sampled at 1 kHz were recorded on a Neurofax EEG-1200 system (Nihon Koden, Japan). All experimental task data were processed off-line using EEGLAB and custom routines programmed in MATLAB (MathWorks, MA, USA). Recordings were down-sampled to 500 Hz and band-passed filtered between 0.1 to 195 Hz using a zero-phase lag filter (FIR). Power line 50 Hz noise and its harmonics were notch-filtered. All the contacts in the white matter were removed. Data were referenced to the common average reference computed from the average of all the clean intracranial electrode channels. The above preprocessing steps were undertaken with a Matlab toolbox EEGLAB^61^.

#### Electrode localization (*Fig 1A*)

Intracranial electrodes were localized on a 3-d MNI brain template using a fusion of pre-operative magnetic resonance (MR) and post-operative computed tomography (CT) scans. We started by co-registering the MR T1 image in subject space to standard MNI space (1mm) using the Advanced Normalisation Tools (ANTs; http://stnava.github.io/ANTs/). This process involves fitting, rigid registration, affine registration, and warping using a diffeomorphic field map (SyN algorithm). Next, we used the affine transformation matrix from this co-registration to register the CT image to standard MNI space. The co-registered and normalized CT and MR images were then visually inspected for the anatomical location of the electrode channels (Fig S1). The localization of the electrodes in MNI space was corroborated with the implantation maps generated from the participant’s own MRI by the epileptologist to clinically localize seizures. The electrode channels were also assessed on the Harvard-Oxford anatomical atlas ^62^ using a Matlab toolbox Brainstorm^63^ to confirm their location in the anatomical regions of interest. The electrode channels of one patient are mapped in MNI for illustrative purposes using BrainNet Viewer^64^ (Fig 2A).

#### Experimental paradigm

Participants were shown the short film *The Butterfly Circus* ^65^, an independent production that has been utilized as a stimulus in prior fMRI research^29,66^. The film was displayed on a 14-inch laptop LCD screen positioned approximately 60 cm from the participants. The movie stimulus was delivered using Presentation® software (Version 18.0, Neurobehavioral Systems, Inc., Berkeley, CA, www.neurobs.com).

#### Data segmentation

To examine dynamic integration and segregation properties, we segmented the continuous iEEG recordings into non-overlapping 5-second windows. This window length was selected to ensure a sufficient number of R-peaks for robust mean HR calculations within each window. Additionally, HR rhythms vary across individuals and are influenced by the emotional engagement elicited by the movie, leading to asynchronous and variable responses. A 5-second window was thus justified to capture dynamic fluctuations while maintaining temporal resolution, enabling reliable analysis of HR-brain network relationships without compromising the ability to track rapid changes in network dynamics.

#### High frequency broadband (HFB) activity

We quantified HFB activity as previously reported^67^. Briefly, time series for each channel was filtered between 70 and 140 Hz using sequential band-pass windows of 10 Hz (i.e., 70– 80, 80–90, 90–100…130-140), via a two-way zero-phase lag FIR filter (EEGLAB). The amplitude (envelope) of each narrow band signal was then calculated from the Hilbert transform. HFB activity is a validated measure of BOLD correlation. Infra-slow LFP activity has also been shown to be reflective of BOLD signals^68^, however, here we did not focus on this due to Nihon Kohden’s sEEG amplifiers typically applying a hardware high-pass filter (HPF) around 0.1–.5 Hz, preventing true DC recordings and attenuating infraslow (<0.5 Hz) signals.

#### Graph measures

In each window for each patient, functional connectivity matrices were constructed by computing pairwise Pearson correlations between regional HFBA. These matrices were binarized and thresholded at a correlation value of 0.2 to retain significant connections while minimizing spurious edges. Integration and segregation metrics were calculated using the Brain Connectivity Toolbox (BCT)^69^. Global efficiency, a measure of integration, was computed to quantify the efficiency of information transfer across the network. Modularity, a measure of segregation, was used to assess the degree of network compartmentalisation into distinct communities. Both metrics were computed for each window, yielding time series of integration and segregation dynamics. We also converted the correlation values to absolute values to take into account the broadness of communication as undertaken in previous studies^70^. Patient-specific Pearson correlations between integration-HR and segregation-HR dynamics were averaged to obtain group-level correlations. Statistical significance was assessed using non-parametric permutation testing (5000 iterations) to generate null distributions.

#### Heart rate

For each patient, ECG was also recorded along with their iEEG. ECG recordings were low-pass filtered (20Hz) and segmented into 5 second epochs. Processed through QRStool^71^ to identify the R peaks, which were visually inspected for misidentified or missed R peaks. Interbeat interval was then calculated from the identified R peaks and converted to HR (60/IBI).

#### Movie features

We extracted movie features—audio, luminance, facial emotions, and visual salience—to study their links to integration, segregation, and HR. Audio dynamics were captured using the Hilbert transform. Luminance and salience were computed using the Pliers toolbox^72,73^ luminance defined as mean pixel intensity and salience derived from computer vision algorithms^74^. Facial emotions were analyzed using the Microsoft API emotion recognition tool ^75^, which detects faces and scores nine emotions (anger, contempt, disgust, fear, happiness, neutral, sadness, surprise). Frames without faces were assigned a score of zero. Emotion scores were summed for each frame. The mean of the above features was computed corresponding to the segmentation of iEEG data. Pearson’s correlation coefficients were computed to find associations between integration, segregation and HR dynamics with that of movie features. False discovery rate was used to control for multiple comparisons at p<.05.

#### Permutation testing

To assess statistically whether the dynamic graph measures were associated with HR, we employed non-parametric permutation testing. We first calculated for each subject the Pearson correlation coefficient between the two corresponding signals of interest. The correlation values were then averaged for all the subjects to obtain a group mean correlation value. To statistically test the significance of the obtained group mean correlation value, we applied non-parametric permutation tests with a null model validated in previous papers^28,53^. Specifically, data from one modality was randomly but circularly shifted relative to the other, before computing the group mean correlation of the null. Five thousand realisations were used to generate the null distribution.

### fMRI data and analyses

Details of the fMRI dataset and preprocessing can be found in a previous publication ^29^. Briefly, 21 healthy participants (11 females and 10 males; all right-handed; aged 21–31 years, with a mean age of 27 ± 2.7 years) watched the same movie, The Butterfly Circus, in the MRI scanner. Three participants were excluded due to excessive head motion, resulting in 18 participants.

Data were acquired using a Siemens TIM Trio scanner with a 12-channel head coil. Functional images were acquired using a gradient-echo EPI sequence (TR = 2200 ms, resolution = 3 mm³). During rest, 220 volumes were collected, while 535 volumes were acquired during movie viewing. A high-resolution T1-weighted structural image (1 mm³) was also obtained. Concurrent physiological recordings included ECG, captured with a BrainAmp MR-compatible amplifier (Brain Products GmbH) at 5000 Hz. Preprocessing was conducted using fMRIPrep 1.1.559 (based on Nipype). Structural images were corrected for intensity non-uniformity and normalized to MNI space. Tissue segmentation (CSF, WM, GM) was performed. Functional images underwent slice-time and motion correction, co-registration to structural images, normalization to MNI space, and smoothing (6 mm Gaussian kernel). ICA-AROMA was applied for denoising. Confound regressors (WM and CSF signals) were extracted, and temporal preprocessing (0.01–0.15 Hz bandpass filtering, WM/CSF regression) was performed using Nilearn. HR data were processed using FMRIB FastR for heartbeat detection, and HR/HRV metrics were derived using the Tapas IO Toolbox^76^. These signals were interpolated and downsampled to match fMRI temporal resolution.

### fMRI data analysis

To assess dynamic brain network properties, preprocessed fMRI data were segmented into 24.2-second windows with a 11-second sliding step. For each window, we followed the same method of quantifying integration (global efficiency) and segregation (modularity) from thresholded (.2) connectivity matrices for each patient. Integration and segregation metrics were calculated using the Brain Connectivity Toolbox (BCT)^69^. Both metrics were computed for each window, yielding time series of integration and segregation dynamics. We also converted the correlation values to absolutes to take into account the broadness of communication as undertaken in previous studies^70^. These integration and segregation time series were obtained for each subject. fMRI preprocessing also yielded HR associated with each TR. Corresponding mean HR for each window were then computed for each subject, and similar sliding window analyses employed to get a mean HR time series corresponding to the integration and segregation metrics. Subject-specific Pearson correlations between integration-HR and segregation-HR dynamics were averaged to obtain group-level correlations. Statistical significance was assessed using non-parametric permutation testing (5000 iterations) to generate null distributions as employed for iEEG analyses

#### Physiological Cardiac response function

To validate our results, we also constructed a physiological regressor by convolving heart rate (HR) time series with a canonical cardiac response function (CRF). The CRF was based on the formulation by Chang et al. (2009), modelled as a double gamma function representing the temporal profile of the BOLD response to cardiac input. Capturing both the early positive and delayed negative components of the cardiac-related BOLD fluctuation spanning 12 seconds. 15 seconds also allows optimally capturing cardiac-induced BOLD responses while preserving sensitivity to rapid autonomic fluctuations during movie-watching. This balances hemodynamic response modelling with temporal precision for naturalistic paradigms. We excluded the first 3 time points of the output signal to avoid convolution-induced edge artifacts. Taking the corresponding data points of integration-segregation time series, we used identical permutation-based testing to test the statistical significance.

#### Integration dynamics with Participation Coefficient

We computed the participation coefficient (PC) for each node using movie-viewing fMRI connectivity matrices, which were thresholded.2 correlation coefficient and binarised. For each subject, we applied undirected modularity detection (modularity_und – BCT toolbox) to identify network communities. Node-wise PC was calculated by assessing the distribution of a node’s connections across modules, with PC = 0 if a node had no connections ^69,77^. Integration for each network was defined as the mean PC across nodes, where higher values reflect greater cross-modular integration (e.g., connector hubs). This was computed dynamically for segmented data using sliding steps consistent with previous fMRI analyses. Subject-level integration time series was then used for subsequent statistical analysis of heart rate associations.

### External fMRI data for validation

We used the open-source fMRI dataset made available Emo-FiLM^78^. The Emo-FiLM dataset is a multimodal resource designed for affective neuroscience research, comprising functional magnetic resonance imaging (fMRI) data collected while participants viewed emotionally evocative film clips. The fMRI data were acquired using a 3 Tesla MRI scanner with a standard echo-planar imaging (EPI) sequence. All functional images were acquired with the same multi-band frequency protocol with a TR/TE/flip angle = 1.3 s/30 ms/64°, field of view 210 mm × 210 mm, slice thickness of 2.5 mm giving a voxel size of 2.5 mm × 2.5 mm × 2.5 mm and whole-brain coverage of 54 interleaved slices. Preprocessing of the fMRI data included motion correction, slice-timing correction, spatial normalization to the MNI template, and spatial smoothing with a 6 mm full-width at half-maximum (FWHM) Gaussian kernel. Heart rate (HR) and respiratory volume per time (RVT) regressors were derived from physiological recordings, convolved with response functions, and included as nuisance regressors^78^. The dataset also includes synchronized physiological recordings (HR and respiration), providing a comprehensive multimodal dataset. We extracted BOLD signals using the AAL atlas, yielding 116 time series. To compute dynamic brain network properties, preprocessed fMRI data were segmented into 26-second windows (20 TRs) with a 6.5 s sliding step (5TRs).

We chose the movie with the greatest engagement as reported by the participants: movie name - “Sintel” of 12:02 (mm:ss) in duration with description of “A young woman embarks on a dangerous quest to find her lost friend, a dragon”, Animation / Fantasy. Absorption-85, engagement - 80, engagement – 83. Data from 30 subjects were available.

### Transforming iEEG signals into BOLD signals with Balloon-Windkessel function

To simulate blood-oxygen-level-dependent (BOLD) signals from intracranial EEG (iEEG) data, we employed the Balloon-Windkessel hemodynamic model, which captures neurovascular coupling by linking neuronal activity to changes in cerebral blood flow and metabolism. The iEEG signals, sampled at 500 Hz, were used as input to the model, representing the neural activity driving vascular responses. A system of differential equations governing blood flow, volume, and deoxyhaemoglobin content was solved using MATLAB’s ode45 solver. The model incorporated key physiological parameters, including neural-to-flow coupling (κ = 0.65), feedback regulation (γ = 0.41), resting oxygen extraction fraction (E = 0.4), vessel stiffness (α = 0.32), time constant for volume change (τ = 0.98), and resting blood volume fraction (V = 0.02). The resulting BOLD signal was computed as a nonlinear function of these physiological states, reflecting the dynamic interplay between neural activity and hemodynamic responses. We applied this transformation to simulated and real iEEG data, validating the output against expected hemodynamic responses. This method enables a computational bridge between electrophysiological and hemodynamic measures, facilitating multimodal neuroimaging analysis. After transformation, we computed graph properties on 25 seconds of windows with a sliding step of 10seconds comparable to the fMRI analyses. Statistical analyses were employed as described above.

## Data and code availability

The data for this project were acquired from patients undergoing clinical care and consenting to additional research protocols. Researchers wishing to access these data will require local ethics approval and a data sharing agreement with Local ethics at QIMR Berghofer and Mater Hospitals Brisbane. Open-source toolboxes have been used for analyses of this study. Specific toolboxes and code are made available on GitHub: https://github.com/srbsonkusare/Integration_segregation.

## Acknowledgements

M.B and R.P. were supported by the National Health and Medical Research Council (APP2008612). M.B. acknowledges funding support of the Advanced Queensland Innovation Partnership (AQIP12316-17RD2). We also acknowledge the QIMR Berghofer Clinician Research Collaboration Grant Award 6626. The authors acknowledge the facilities and scientific and technical assistance of the National Imaging Facility (NIF), a National Collaborative Research Infrastructure Strategy (NCRIS) capability, at the Hunter Medical Research Institute Imaging Centre, University of Newcastle. The funding bodies had no role in study design, analysis, decision to publish, or preparation of the manuscript.

## Author contributions

S.S.: conceptualization, task designing, acquisition, data curation, investigation, software, validation, and formal analysis, visualization, writing - original draft, and writing - review & editing. K.Y..: investigation, software, validation, and formal analysis, writing - review & editing. L.H.: writing - review & editing. R.P.: formal analysis, writing - review & editing. J.M.: curation, writing - review & editing. S.D.: data acquisition, curation, writing - review & editing. J.S.: writing - review & editing. M.B.: conceptualization, funding acquisition, project administration, resources, supervision, writing - review & editing.

