## Supplementary Materials for "Brain-heart coupling shapes large scale brain dynamics"

Sonkusare et al.

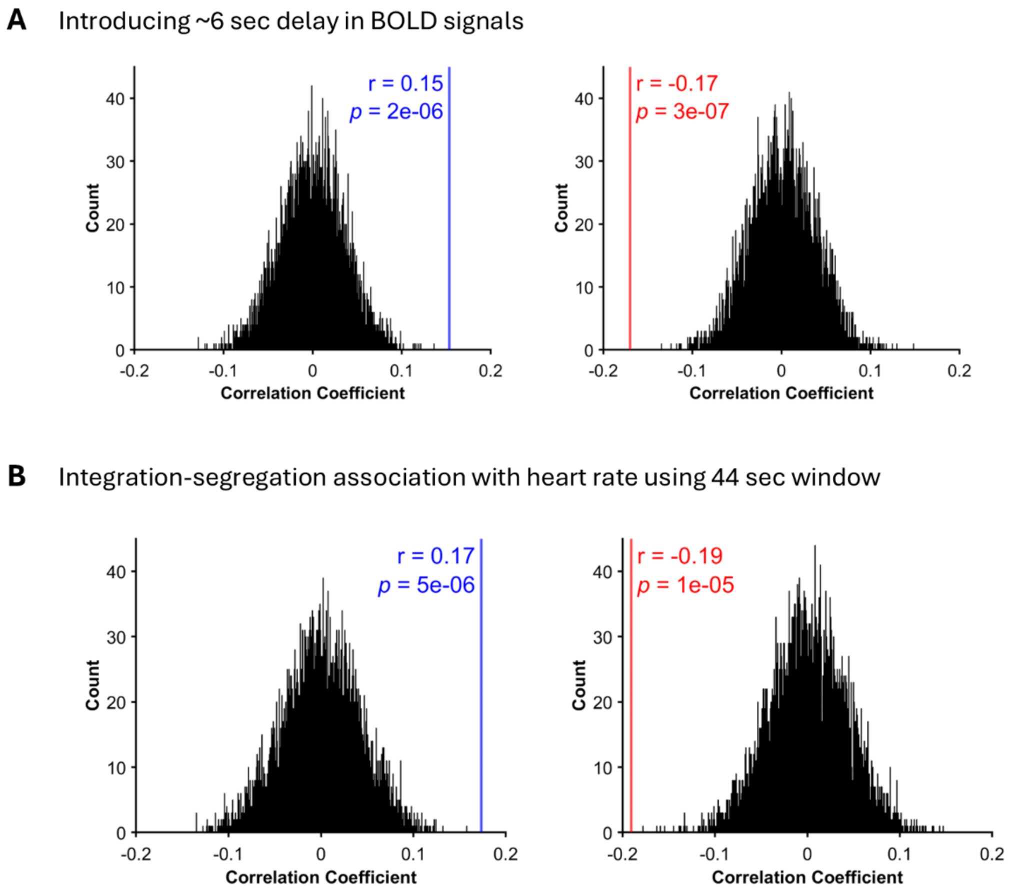


**Figure S1. Replication of BOLD–heart rate association adding 6 second delay and with longer window duration. (A)** Introducing a 6.6 s delay (3 TRs) in BOLD for association with fMRI graph measures of segregation (left) and integration (right) replicates the original findings observed without a delay. Null distribution derived from 5,000 permutations confirms the statistical significance of a positive correlation between segregation and heart rate and a negative correlation between integration and heart rate. **(B)** Extending the window duration to 44 s (20 TRs) reproduces the original pattern of results, consistent with findings using a 24.2 s (11 TRs) window.

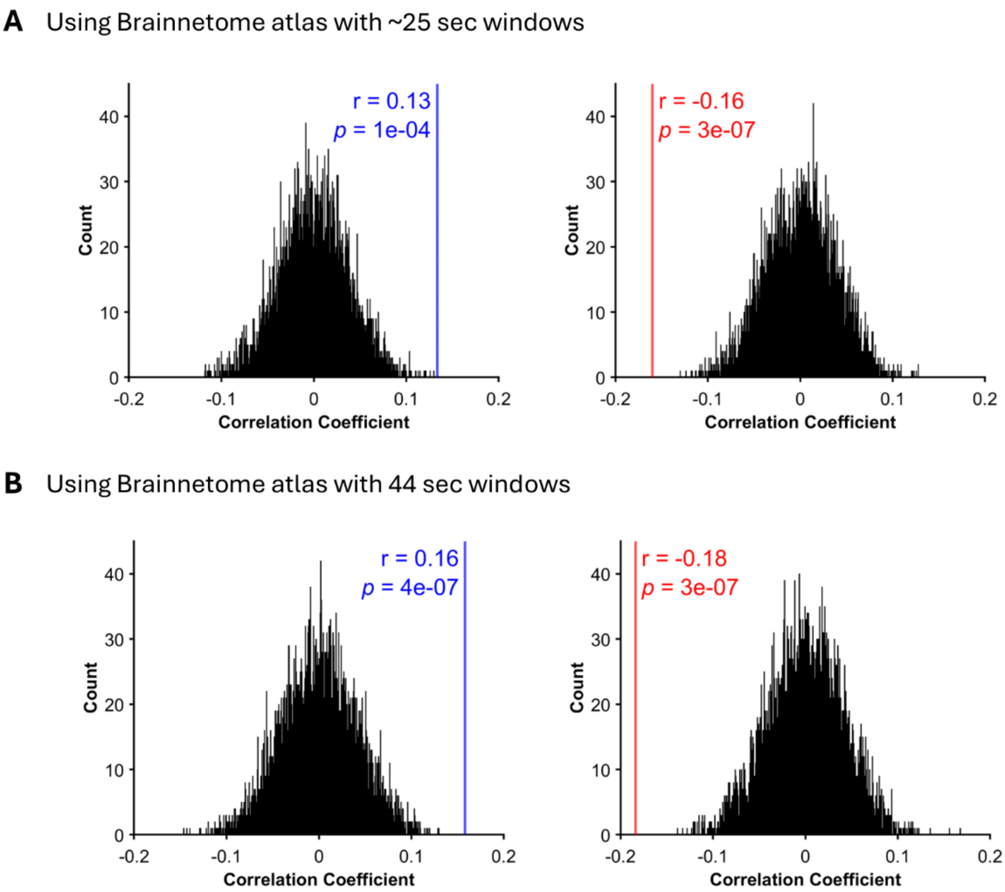


**Figure S2. Replication of BOLD–heart rate association using the Brainnetome atlas. (A)** Using the Brainnetome atlas (274 parcels: 210 cortical, 36 subcortical, 28 cerebellar regions) replicates the original findings of a positive correlation between segregation and heart rate (left) and a negative correlation between integration and heart rate (right) when using a 24.2 s (11 TRs) window. Null distribution derived from 5,000 permutations. **(B)** Obtaining integration and segregation time series with the Brainnetome atlas using a 44 s (20 TRs) window similarly reproduces the original pattern of positive correlation between segregation and heart rate (left) and a negative correlation between integration and heart rate (right), observed with the 24.2 s (11 TRs) window.

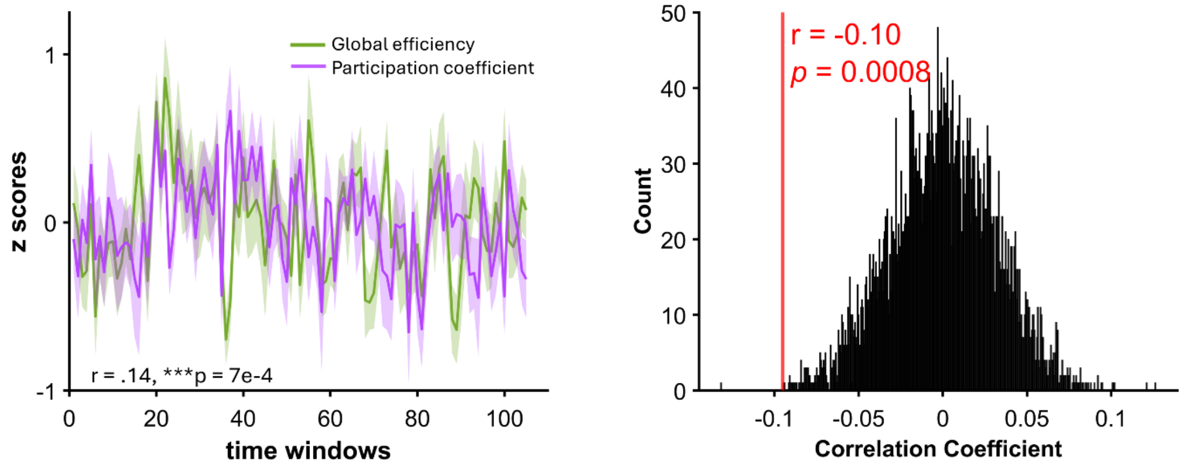


**Figure S3. Integration dynamics with participation coefficient.** Significant similarity was found between integration dynamics obtained via global efficiency and via participation coefficient (left). Using integration dynamics from participation coefficient also showed similar negative association with heart rate dynamics. Null distribution obtained via permutation testing.

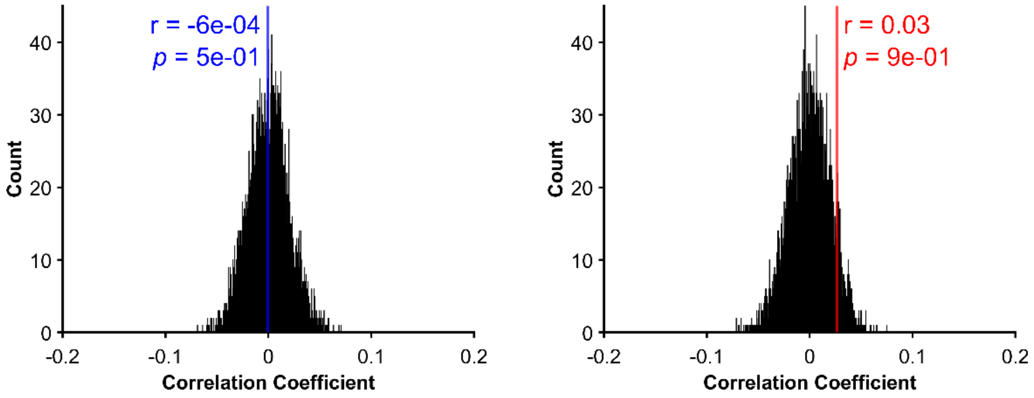


**Figure S4. Association of graph properties on iEEG alpha activity and heart rate.** Alpha frequency activity (8–12 Hz) has been shown to be negatively correlated with BOLD signals, reflecting an inverse relationship with fMRI-derived neural activity. We applied the same analysis used for iEEG high-frequency activity to examine the association between heart rate and graph properties of segregation (left) and integration (right), with iEEG data bandpass filtered in the alpha range. Mean correlation values showed no statistical significance when tested against a null distribution generated from 5,000 permutations.
